# MateR: a novel framework for computing the usefulness criterion and applying genomic mating

**DOI:** 10.1101/2025.09.24.678244

**Authors:** Javier Fernández-González, Seifelden M. Metwally, Julio Isidro y Sánchez

## Abstract

Genomic mating uses genome-wide information to design crosses that maximize genetic gain while managing diversity. Expected gain is often predicted through the usefulness criterion, which depends on family means and variances. However, existing equations mix incompatible parametrizations when considering dominance effects. Furthermore, diversity control is often tuned with metrics that lack a direct link to loss of additive variation and long-term gain. We derived equations that compute family mean and within-family variance consistently under breeding and genotypic parametrizations by computing locus-specific values using genotypic frequencies and propagating them to the entire genome through linkage disequilibrium covariances. We also developed a diversity metric that estimates the proportion of additive standard deviation lost and integrated both advances into the MateR software. We evaluated performance in simulated diploid and autotetraploid crop populations across multiple breeding schemes and against existing tools. The new equations predicted family means and variances with near-perfect accuracy when true QTL effects were known. With estimated marker effects, correlations were roughly 0.55–0.90 for family mean and about 0.25 for within-family standard deviation. The diversity metric matched the expected loss of additive standard deviation under random sampling and tracked loss of genic variance under selection. This framework unifies prediction of cross usefulness under dominance and supplies an interpretable diversity control directly tied to long-term gain. Implemented in MateR, it applies to diploids and autopolyploids and accommodates common breeding program constraints, including hybrid schemes and testers.

## Introduction

Breeding for quantitative traits relies on crossing elite parents to generate new diversity and select superior genotypes. Optimizing a mating plan requires predicting the outcome of every possible cross, which has been made possible by genomic prediction (GP) (Meuwissen *et al*. 2001). This optimization, often termed genomic mating or optimal mate allocation (Akdemir and Sánchez 2016; Gorjanc and Hickey 2018), aims to maximize genetic gain as described by the breeder’s equation:

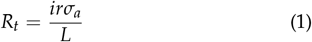

Where *R*_*t*_ is the response to selection per unit of time, *i* is selection intensity, *r* is prediction accuracy, *σ*_*a*_ is the additive genetic standard deviation, and *L* is the generation interval (Lynch *et al*. 1998). In genomic mating, *r* and *L* are typically held fixed, whereas the mating plan can alter *i* and *σ*_*a*_. Prioritizing only the best parents inflates *i* in the short term but rapidly depletes *σ*_*a*_ across cycles. Thus, early implementations (Kinghorn 2011; Akdemir and Sánchez 2016; Gorjanc and Hickey 2018) balanced two objectives, maximizing the average merit of selected parents (strongly related to *i*) and maintaining genomic diversity to limit the loss of *σ*_*a*_. This is a relatively simple and effective strategy (Endelman 2025) that nonetheless overlooks parental complementarity at the cross level.

Two well-known features underscore this limitation. First, within-family variance affects the chance of sampling exceptional progeny. To account for this, the *usefulness criterion* (Schnell and Utz 1975; Zhong and Jannink 2007; Lehermeier *et al*. 2017b) was developed to integrate family mean and variance. Second, dominance can induce transgressive segregation and heterosis (Falconer 1989), elevating progeny performance beyond the mid-parent value. Historically, these effects could only be measured by phenotyping the progeny, which would defeat the point of optimizing the mating plan. In contrast, nowadays GP allows to predict both quantities for any pair of genotyped parents (Lehermeier *et al*. 2017b), enabling refined genomic mating. Akdemir and Sánchez (2016) incorporated within-family variance, and recent work extended usefulness to incorporate dominance effects (Wolfe *et al*. 2021; Peixoto *et al*. 2025). However, these efforts unintentionally mixed distinct GP parametrizations, which compromises predictions.

The commonly used *breeding parametrization* (Vitezica *et al*. 2013) decomposes total genotypic value into breeding values and dominance deviations in the absence of epistasis. The breeding value of a genotype is its expected progeny mean when mated at random to a reference population, and includes all additive effects plus the expected dominance. The dominance deviations are simply the difference between the total genotypic value and the breeding values. Critically, breeding values and dominance deviations are independent, which is a key assumption made by Wolfe *et al*. (2021) and Peixoto *et al*. (2025) when computing family variance. In contrast, their computation of family mean followed Falconer (1989) (see Equation 2). This approach assumes the *genotypic parametrization*, which decomposes total genotypic value into strictly additive and strictly dominance components that are non-independent from each other (Vitezica *et al*. 2013). Consequently, if one adopts the breeding parametrization, the mean is miscomputed; and if one adopts the genotypic parametrization, the variance assumptions fail. Our first objective is, therefore, to derive equations for family mean and variance that remain accurate under both parametrizations, enabling correct usefulness under dominance.

Our second objective addresses diversity control. Genomic mating balances usefulness against the expected depletion of *σ*_*a*_, typically called inbreeding control. Crucially, the importance given to each factor is controlled by a user-defined hyperparameter that has to be tuned (Akdemir and Sánchez 2016). We argue that an easy-to-interpret diversity metric makes the tuning much easier and intuitive. Common approaches use kinship between selected parents (Akdemir and Sánchez 2016; Peixoto *et al*. 2025) or kinship-based angles (Kinghorn 2011; Gorjanc and Hickey 2018), which work well but are difficult to interpret and tune, as it is very difficult to link them to specific values of *σ*_*a*_ depletion and long-term gain. The classical inbreeding rate (Wray and Thompson 1990; Toro and Perez-Enciso 1990; Wray and Goddard 1994; Meuwissen 1997; Grundy *et al*. 1998; Endelman 2025) is interpretable but hard to estimate robustly from markers without pedigrees that separate identity by descent (IBD) from identity by state (IBS) (Wang 2014; Endelman 2024). Furthermore, it does not directly quantify the expected reduction in *σ*_*a*_ per cycle, and therefore it is difficult to link it to long-term gain potential, complicating the tuning. We therefore aim to develop a diversity metric that is directly interpretable as the anticipated reduction in additive standard deviation for a given mating plan.

Finally, our third objective is to implement these developments in a new R package called `MateR`, delivering genomic mating with (i) parametrization-consistent usefulness under dominance and (ii) an interpretable handle on diversity loss. We emphasize versatility, supporting autopolyploid species, as well as clonal, self-pollinated, and hybrid crops; together with animal breeding.

### Mathematical Development

For clarity, we present derivations under the *genotypic parametrization* in this section. All results hold under the *breeding parametrization* provided the corresponding marker scores and effects are used.

#### Family mean

The family mean is often computed as the mid-parent value (valid only under additivity) or using Equation 14.6 in Falconer (1989):

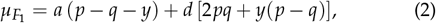

which gives the expected genotypic value of the *F*_1_ from two distinct parental populations. This expression is restricted to *F*_1_ and the genotypic parametrization, and in our tests it did not reproduce the correct expectations (Supplementary File 1). We therefore propose a more general and accurate method based on genotypic frequencies.

Let *F* denote the family generated by crossing parents *P*_*k*_ and *P*_*l*_. Consider locus *j* with alleles *A* and *a*, and additive and dominance effects *a*_*j*_ and *d*_*j*_. Under diploidy (or disomic inheritance), the additive marker score *M*[*i, j*] ∈ {0, 1, 2} for individual *i* is encoded as the count of allele *a* (*AA* = 0, *Aa* = 1, *aa* = 2), and the dominance score *W*[*i, j*] ∈ {0, 1} as a heterozygosity indicator (*AA* = 0, *Aa* = 1, *aa* = 0). Let *PAA*_*F,j*_, *PAa*_*F,j*_, and *Paa*_*F,j*_ be the genotypic frequencies in *F* at locus *j* (summing to 1). The locus-specific family mean can be simply calculated by applying the definition of arithmetic mean:

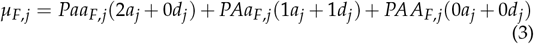

This requires only the marker effects and the genotypic frequencies in the target family, which can be computed for *F*_1_, doubled haploids, arbitrary generations of selfing, backcrosses, and tester designs (Appendix 1). Thus, Equation 3 can be easily applied to any parametrization (given the corresponding scores/effects) and any generation, solving the limits of Equation 2. Furthermore, it also extends to autopolyploids by replacing the diploid encodings with their polysomic counterparts (Appendix 1).

Equation 3 refers to a single locus, but it can be easily extended to the entire genome. The total genotypic value (*g*_*i*_) of an individual *i* is computed by summing over the additive and dominance effects across all *n*_*j*_ loci:

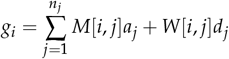

Accordingly, the expected genotypic value of a family *F* is:

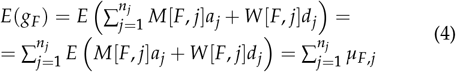

#### Within-family variance

Using the same notation as above and the definition of variance, the per-locus within-family variance for family *F* at locus *j* is

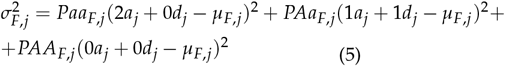

Where 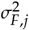 is the within-family variance for locus *j* and *µ*_*F,j*_ is calculated from Equation 3. Extensions to autopolyploids follow by replacing the diploid genotype encodings with their polysomic counterparts, see Appendix 1.

Across the genome, the variance of the sum of locus contributions includes both locus variances and between-loci covariances:

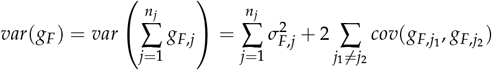

In the literature, this has been computed as (Lehermeier *et al*. 2017a,b; Wolfe *et al*. 2021; Peixoto *et al*. 2025):

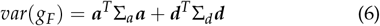

where ***a*** and ***d*** are vectors of additive and dominance marker effects, and Σ_*a*_ and Σ_*d*_ are the covariance matrices between the additive and dominance marker scores (doses). This works fine for the breeding parametrization, which assumes that additive and dominance effects are independent. However, for the genotypic parametrization, we would need to consider their correlation. Therefore, we propose an alternative formulation where we instead express the genome-wide variance via single-locus standard deviations and their correlations.

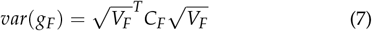

Where 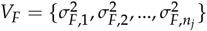 are locus-specific variances from Equation 5 and *C*_*F*_ is the correlation matrix between loci, i.e., it reflects the linkage disequilibrium (LD). Importantly, *C*_*F*_ is the correlation of the genotypic effects of the loci, not just of the marker scores. This formulation implicitly accommodates any correlation between additive and dominance effects present under the genotypic parametrization. Computing *C*_*F*_ is complicated, and we have developed three options, ranging from very simplistic to very realistic:

1. **Independent loci**. Assume *C*_*F*_ = *I*, where *I* is an identity matrix with the needed dimensions. This results in 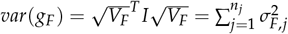 and is equivalent to the approach in Akdemir and Sánchez (2016).
2. **Parental-LD propagation**. *C*_*F*_ can be computed as:

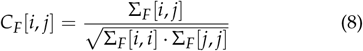 Where Σ_*F*_ refers to the covariance matrix across loci for the genotypic values in family *F*. Σ_*F*_ can be calculated as:

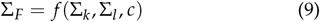 Where Σ_*k*_ and Σ_*l*_ denote the covariance matrices of the parental lines, and *c* is the matrix of recombination frequencies. The function in Eq. 9 depends on the breeding scenario. For example, when the parents are not fully homozygous and the *F* family corresponds to the F1 generation,

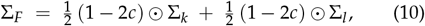 Where ⊙ denotes the Hadamard (elementwise) product. See Lehermeier *et al*. (2017b) for expressions of Σ_*F*_ for doubled haploids and for any number of selfing generations. The last step is to compute Σ_*k*_ and Σ_*l*_ to evaluate the variance. The same procedure applies to both parents; for brevity, we illustrate it for parent *P*_*k*_:
  - Define a matrix of genotypic effects for the population from which the parent *P*_*k*_ was taken:

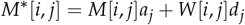
  - Center columns to obtain *Z*^*∗*^.
  - Compute: 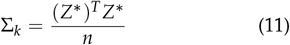 Where *n* is the number of individuals in the population. In summary, within-family variance can be calculated by applying equations 5, 11, 9, 8 and 7 in that order.
3. **Compute LD for each individual parent:** In the previous scenario, we assumed that LD in each parent equals LD in its source population. A more realistic approach is to compute LD for specific parents, which requires phased genotypes (trivial for fully homozygous lines but rarely available otherwise). Let *H*_*k*_ be the *ϕ* × *n*_*j*_ haplotype matrix for parent *P*_*k*_, where *ϕ* is the degree of ploidy, rows represent chromosomes and columns represent markers. Each cell is coded as 0 (reference allele) or 1 (alternative allele). We compute allele frequencies in the family as the average from both parents:

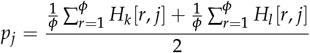 To compute the covariance matrix of additive values across loci, we define the following:
  - Centered matrix of phased additive effects:

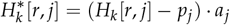
  - Additive covariance matrix:

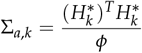

It is important to note that, in this scenario, the covariance matrix Σ_*a,k*_ does not reflect any dominance. The reason for this is that the phased information in *H*_*k*_ is purely additive, and there is no way to get phased dominance marker scores, as the dominance in a locus is not assigned to a specific chromosome, but it is rather the outcome of the interaction among all of them. Next, it is also important to highlight that *p*_*j*_ has to be the allele frequency in the family. The other possibility would be setting *p*_*j*_ as the allele frequency in parent *P*_*k*_ (Wolfe *et al*. 2021), but this would cause all homozygous loci to become monomorphic. As a result, their variance and covariances would become zero, which is not realistic.

Next, we can find the additive covariance between loci in the family from Σ_*a,k*_, Σ_*a,l*_ and *c* using Equation 9. It can be subsequently converted into a correlation matrix *C*_*a,F*_ using Equation 8.

Finally, we need to include dominance. To that end, the first step is finding the additive and dominance variances within each locus 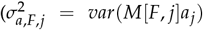 and 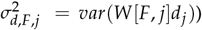. They can be obtained using Equation 5 for total variance and setting *d*_*j*_ = 0 or *a*_*j*_ = 0 to isolate additive or dominance variance respectively. Finding them allows us to calculate the correlation between additive and dominance effects. For any locus *j* in family *F*, the genotypic variance is:

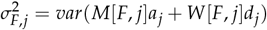

Expanding this:

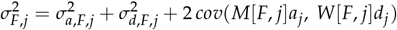

Solving for the covariance and correlation gives:

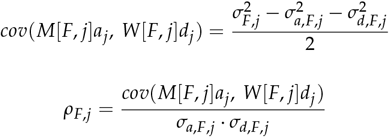

These expressions allow us to obtain the within-family genotypic variance using path analysis Wright (1934). For instance, following Figure 1, we can get the correlation between the dominance effects in loci 1 and 2 as *ρ*_*F*,1_*C*_*a,F*_ [1, 2]*ρ*_*F*,2_.

**Figure 1.**
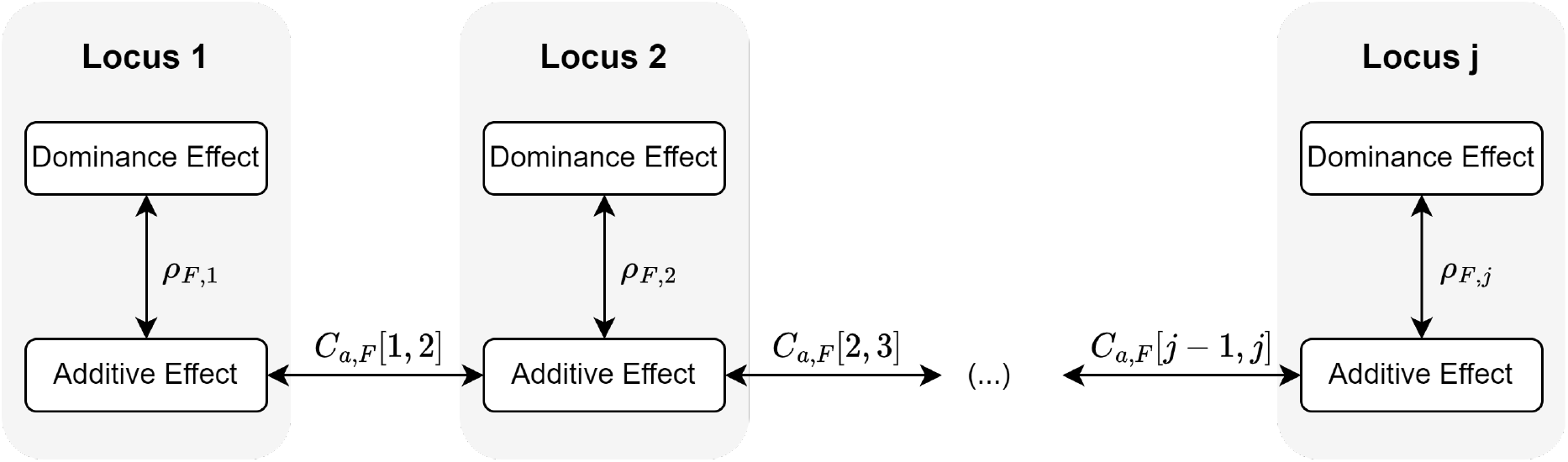
Scheme of the path analysis for the additive and dominance effects in each locus for family *F*.

Using Figure 1, and the bilinearity of covariance, we can compute the total within-family genetic variance by summing all additive and dominance covariance components:

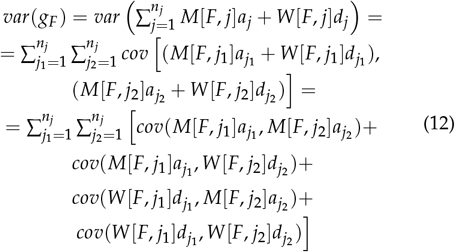

Each of the terms above can be computed using *C*_*a,F*_ and *ρ*_*F,j*_ as follows:

#### Additive *×* additive covariances

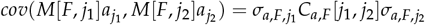

#### Additive *×* dominance covariances

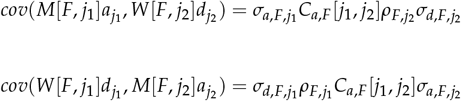

#### Dominance *×* dominance covariances

If *j*_1_*≠ j*_2_:

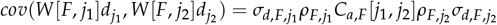

If *j*_1_ = *j*_2_:

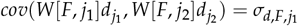

This distinction ensures that when *j*_1_ = *j*_2_, we do not incorrectly scale the variance by 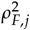, since there is a direct self-connection in the path diagram (no indirect dependency).

In summary, within-family variance using phased genotypes can be computed as:

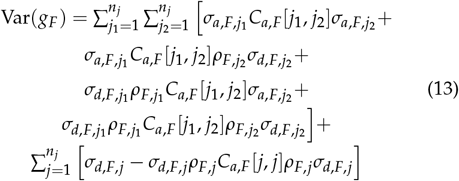

All equations above have been developed for single-trait analysis. Extension to multi-trait is available in Appendix 2.

#### Proportion of additive standard deviation lost

Controlling genetic diversity is central to genomic mating. A simple, interpretable metric helps tune constraints and makes explicit how selection trades short-term response for long-term potential. Although inbreeding rate is commonly used, its accurate estimation typically relies on pedigrees (Wang 2014; Endelman 2025) and it does not directly translate into the expected reduction in genetic gain. We therefore define the *proportion of additive standard deviation lost* (*PropSD*), which uses only genomic information and links directly to long-term gain because the breeder’s equation scales linearly with additive standard deviation.

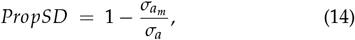

where *σ*_*a*_ and 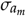 are the additive standard deviations for the available parental pool and for the selected parents of the mating plan. A naive approach would compute *σ*_*a*_ and 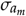 as the sample standard deviations of their genomic estimated additive values. However, this requires marker effects, which are extremely difficult to accurately estimate in quantitative traits (Rice *et al*. 2008; Korte and Farlow 2013; Ramstein *et al*. 2019; Clauw *et al*. 2024). Thus, in the naive approach, it is likely that small effects are wrongly assigned to some important positions, minimizing their impact on the value of *PropSD* and removing any penalization for the fixation of undesired alleles in these loci. To avoid dependence on marker effects, we assume 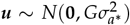, where *G* is a genomic relationship matrix with associated variance com-ponent 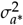 and ***u*** is a vector of additive values. The expected sample standard deviation of ***u*** from a random sample can be written as:

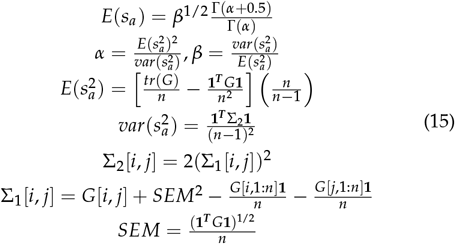

Where Γ(·) refers to the gamma function, *tr*(·) refers to the trace of a matrix, and *n* is the number of rows or columns in *G*. A detailed mathematical derivation of Equation 15 is available in Appendix 3.

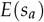 can be used as an estimator for *σ*_*a*_. We can similarly estimate 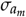 from 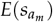, computed using Equation 15 with *G*_*m*_ instead of *G*. This allows us to estimate *PropSD* as:

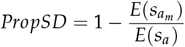

While *PropSD* is the primary method for controlling diversity loss in MateR, inbreeding rate and standard error of the mean are also supported. More details in Appendices 4 and 5.

#### Objective function for the optimization

For each family *F*, we define usefulness as’

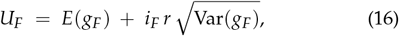

where *E*(*g*_*F*_) and Var(*g*_*F*_) are obtained from Equations 4 and 7, *r* is the prediction accuracy associated to the marker effects employed, and *i*_*F*_ is the within-family selection intensity, which can be computed as a function of the proportion of individuals selected within the family. In the optimization process, it is often desired to replicate a cross, i.e., mating two parents more than once to generate more offspring, which increases *i*_*F*_. This makes the value of *i*_*F*_ dependent on the mating plan, which results in a non-convex optimization problem.

For a mating plan *m* producing *n*_*F*_ families, we define fitness as the average usefulness across all families:

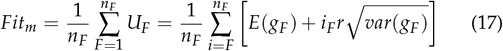

The optimization problem is then:

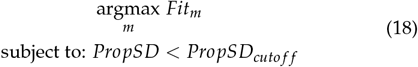

where *PropSD* follows Equation 14 and *PropSD*_cutoff_ is a user-specified threshold on the acceptable loss of *σ*_*a*_.

## Materials and methods

### Optimization algorithm

Given the objective in Equation 18, we maximize it with `TrainSel` (Akdemir *et al*. 2021b), a hybrid heuristic that blends a genetic algorithm with simulated annealing (Akdemir *et al*. 2021a). We ruled out convex optimization and integer programming because the fitness landscape is non-convex and includes discrete replication decisions. The optimization procedure is:

1. Pre-compute family mean and within-family variance for all possible crosses.
2. Initialize the TrainSel heuristic by creating random mating plans.
3. For each mating plan considered by TrainSel:
  3.1. Compute *PropSD*. If exceeds the user threshold, set a very small value to *Fit*_*m*_ and jump to step 3.4.
  3.2. Similarly enforce any other constraints.
  3.3. Compute *Fit*_*m*_ from Equation 17.
  3.4. The mating plans with highest *Fit*_*m*_ are classified as elite and the rest are discarded. Mating plans with lower *Fit*_*m*_ may also be accepted during simulated annealing steps.
  3.5. If convergence or maximum number of iterations is reached, stop. Otherwise, create new mating plans by applying mutation and crossover operators to the elite solutions and go back to step 3.

MateR supports numerous constraints for step 3.2. To maximize versatility, MateR differentiates between a family and a cross. A family is the set of offspring obtained by crossing two parents. However, these two parents may be crossed more than once. The more crosses are performed for a given family, the larger the number of offspring for that family. This framework allows including the following constraints:

- Fix the total number of crosses (program-level progeny budget).
- Enable or disable repeated crosses per family. If it is disabled, there is always exactly one cross per family.
- Fix the number of families to be represented (optional).
- Impose parent-level bounds: minimum and/or maximum number of crosses per parent (optional).
- Specify allowed and forbidden crosses between any pair of parents (optional).
- Include a set of testers (optional); the algorithm then optimizes parental matings to maximize the performance of the hybrids resulting from crossing the progeny with the testers, accounting for general combining ability (GCA) and specific combining ability (SCA).

To our knowledge, `MateR` is the first genomic mating software to natively optimize mating plans conditional on a user-specified tester set, a feature that is particularly valuable for hybrid breeding.

A detailed guide to MateR usage is available in https://github.com/TheRocinante-lab/MateR

### Simulated datasets

We simulated one diploid dataset and one autotetraploid dataset. These serve as the `MateR` example data and were used to validate the proposed equations.

Each dataset contained 1,000 loci distributed across 8 chromosomes of 150 centimorgans (cM) each. For every chromosome, locus allele frequencies were sampled independently. We then imposed a target autocorrelation of 0.95 between adjacent loci along the genetic map. When the maximum feasible correlation between two loci was smaller than 0.95 due to their allele frequencies, the target was reduced to the largest value compatible with those frequencies. Founder haplotypes were sampled iteratively to satisfy the frequency and LD targets, yielding populations with strong linkage disequilibrium, by design, to rigorously probe LD-sensitive variance formulas.

In the diploid population, we created two subpopulations (heterotic-pool analogs), each with 50 fully-homozygous lines. For each subpopulation, we first simulated 200 haplotypes, performed 50 random crosses, and converted the resulting progeny to doubled haploids. Allele frequencies and LD patterns were simulated independently for each subpopulation.

In the autotetraploid dataset, we simulated a single subpopulation with 400 founders and carried out 100 random crosses, producing 100 non–fully homozygous genotypes.

Finally, we simulated marker effects for two traits, Yield (YLD) and maturity (MAT). Marker effects followed the genotypic parametrization and were drawn from a multivariate normal distribution with correlation 0.3 between traits.

### Validation of family mean and variance

To validate the accuracy of the predicted family mean and within-family variance, we sampled a random pair of parents, simulated 10^4^ progeny per cross, and took the sample mean and variance of the simulated progeny as the true values for that family. We then compared these values to predictions from `MateR`, and from other genomic mating software, `SimpleMating` (Peixoto *et al*. 2025), `predCrossVar` (Wolfe *et al*. 2021), and `Genomic Mating` (Akdemir and Sánchez 2016). Method names and the corresponding linkage disequilibrium (LD) assumptions are summarized in Table 1. As performance metrics, we computed i) Pearson correlation (*accuracy*) and ii) normalized root mean square error (NRMSE = RMSE divided by the standard deviation of the true values). The entire procedure was repeated for 500 independent crosses.

**Table 1.** Name used for each estimator of family mean and variance, classified by how LD is handled and by the software used. Numbers in brackets refer to the corresponding subsections in the Mathematical Development describing the three variance formulations. For `GenomicMating`, we used only the approaches described in the original publication to avoid redundancy with later updates that are very similar to `SimpleMating`.

| Method | MateR | SimpleMating | predCrossVar | GenomicMating |
| --- | --- | --- | --- | --- |
| Family Mean | MateR | SM | pCV | GM |
| Family Variance, independent loci (1) | MateR_Ind | - | - | GM |
| Family Variance, approximate LD (2) | MateR_Approx | SM_Approx | - | - |
| Family Variance, phased genotypes (3) | MateR_Full | SM_Full | pCV | - |

We evaluated five breeding schemes:

- **DH:** line breeding program with double haploids.
- **RILs:** line breeding program with 3 selfing cycles before yield trials.
- **Hybrid_DH:** hybrid breeding program with double haploids.
- **Hybrid_RILs:** hybrid breeding program with 3 selfing cycles before crossing with testers.
- **Clonal:** clonal breeding program, no selfing cycles.

All schemes were implemented in the diploid population and non-double-haploid programs were also implemented in the autotetraploid population, yielding 8 total scenarios. Not all methods in Table 1 apply to every scenario because some packages support only a subset of breeding schemes.

Within each scenario, we randomly sampled 250 loci as QTL, and defined offspring true genotypic values (TGVs) from the corresponding QTL effects. Predictions of family mean and family variance were produced under four genomic prediction settings:

- **QTL_true**: the true QTL effects were used.
- **QTL_1**: estimated QTL effects were obtained. These effects were different from the true QTL effects, but resulted in perfect predictions of the genome-wide genotypic values of the parental lines (although predictions were no longer perfect for the offspring).
- **QTL_0.7**: we used estimated QTL effects such that genomic estimated genotypic values (GEGVs) of parents had a correlation of 0.7 with TGVs.
- **Markers_0.7**: marker effects were obtained for the 750 non-QTL loci. These marker effects resulted in GEGVs of the parents that presented a correlation of 0.7 with the TGVs.

### Validation of diversity constraint

We tested whether *PropSD* accurately reflects the expected reduction in additive standard deviation. For each of the diploid and autotetraploid datasets, we formed parental samples of size 5, 20, 100, and 200 *with replacement*, repeating each size 20 times (80 samples per dataset). Additive genetic values were modeled as ***u*** ~ *N*(**0, G**), with **G** being the genomic additive relationship matrix of the full parental pool.

Within each parental sample *i*, we drew 10^5^ sets of marker effects. Therefore, for each draw *j*, we had a different vector of additive values for the parental pool (***u***_*j*_). Next, we computed the realized loss of additive variability as

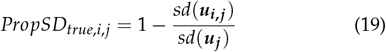

where ***u***_*i,j*_ denotes the vector of additive values for the sampled parents. We then compared the predicted *PropSD*_*i*_ from Eq. 14 with the Monte Carlo average, i.e., 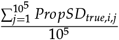.

Because random sampling underestimates the coupling between loci induced by directional selection, we additionally evaluated *PropSD* under selection. Using `MateR`, we optimized mating plans subject to *PropSD*_cutoff_ ∈ {0.001, 0.01, 0.02, 0.03, 0.04, 0.05}, and for each cutoff generated 100 independent sets of marker effects used for optimization. We compared predicted and realized *PropSD* in both ploidy levels, computing realized loss from both (i) the *genetic* standard deviation (captures LD) and (ii) the *genic* standard deviation (excludes LD) to examine Bulmer’s variance depletion under selection (Bulmer 1971).

## Results

Here, we evaluated how predictions for family mean, within-family standard deviation, and *PropSD* agreed with their simulated true values across ploidy levels and scenarios.

### Family mean

Across additive values, all packages performed identically and achieved perfect accuracy when the true QTL effects were known (Figure 2A,B). For genotypic values, `MateR` was superior: it uniquely reached perfect accuracy and was the only software that operated across all scenarios, including the testcross setting in which we predicted the mean of hybrids produced by crossing a family to a fixed tester.

**Figure 2.**
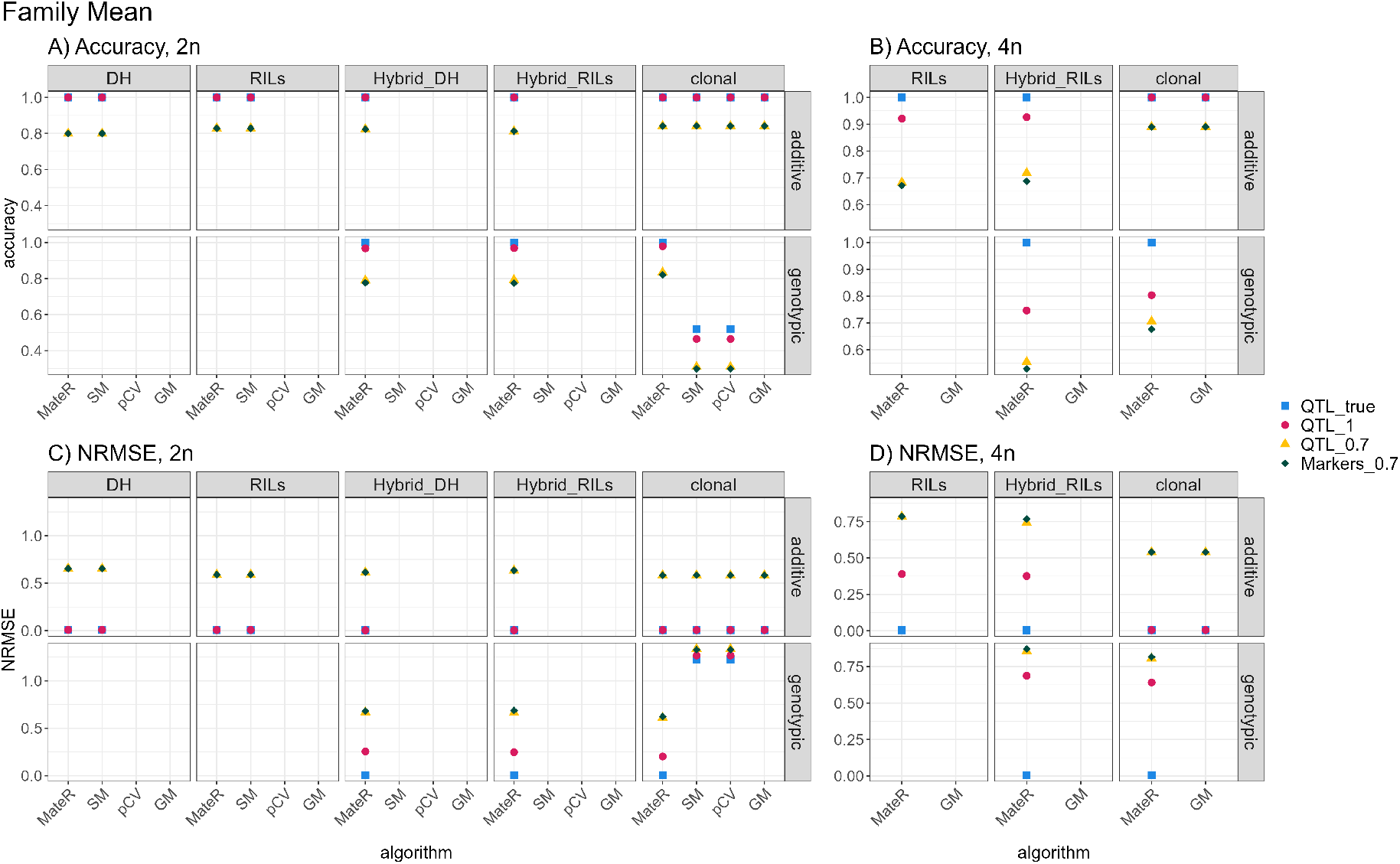
Accuracy (Pearson’s *r*; A–B) and error (NRMSE; C–D) between predicted and true family means across software packages. Panels show diploids (2n; A,C) and autotetraploids (4n; B,D). Within each panel, the top row uses additive values; the bottom row uses genotypic values (additive+dominance). Facet columns are population types (DH, RILs, Hybrid_DH, Hybrid_RILs, clonal). The *x*-axis lists the algorithm used to compute family means (MateR, SM, pCV, GM). Colors/shapes indicate how the effects used for prediction were obtained: 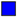 **QTL_true**; true QTL effects; 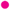 **QTL_1**; estimated QTL effects that exactly reproduce parental genome-wide genotypic values (perfect for parents, not necessarily offspring); 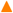 **QTL_0.7**; estimated QTL effects whose parental genomic estimated genotypic values (GEGVs) correlate 0.7 with true genotypic values (TGVs); 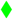 **Markers_0.7**; marker effects fitted on the 750 non-QTL loci producing parental GEGVs with correlation 0.7 to TGVs.

The scenario `QTL_1` used QTL effects distributed differently from the truth while preserving identical genome-wide predictions in the parental population. Its impact on family-mean accuracy and NRMSE was negligible: no effect for additive values in the diploid and clonal autotetraploid programs, and a small negative effect for additive values in the non-clonal autotetraploid scenarios and for genotypic values.

Scenarios `QTL_O.7` and `Markers_O.7` targeted a predictive accuracy in the parental pool of ≈ 0.7. In diploids, family-mean accuracy consistently exceeded this baseline (around 0.8). In autotetraploids, accuracy was less uniform and ranged from 0.55 to 0.90, with higher values in the clonal program and for additive effects.

As expected, NRMSE mirrored these trends (Figure 2C,D). Most importantly, when predicted and true values were perfectly correlated, NRMSE equaled zero, indicating unbiased predictors.

### Within-family standard deviation

`MateR_Full` (phased genotypes as input) was the only method that achieved perfect or near-perfect prediction of within-family standard deviation across all scenarios (Figure 3). `SM_Full` (SimpleMating) matched this performance only in diploids for additive values when parental lines were fully homozy-gous—specifically, the *DH* and *RILs* scenarios in Figure 3A. In the diploid *clonal* scenario of Figure 3A, `MateR_Full`, `SM_Full`, and `pCV` all used phased genotypes, yet only `MateR_Full` translated these inputs into perfect predictions for both additive and genotypic standard deviation.

**Figure 3.**
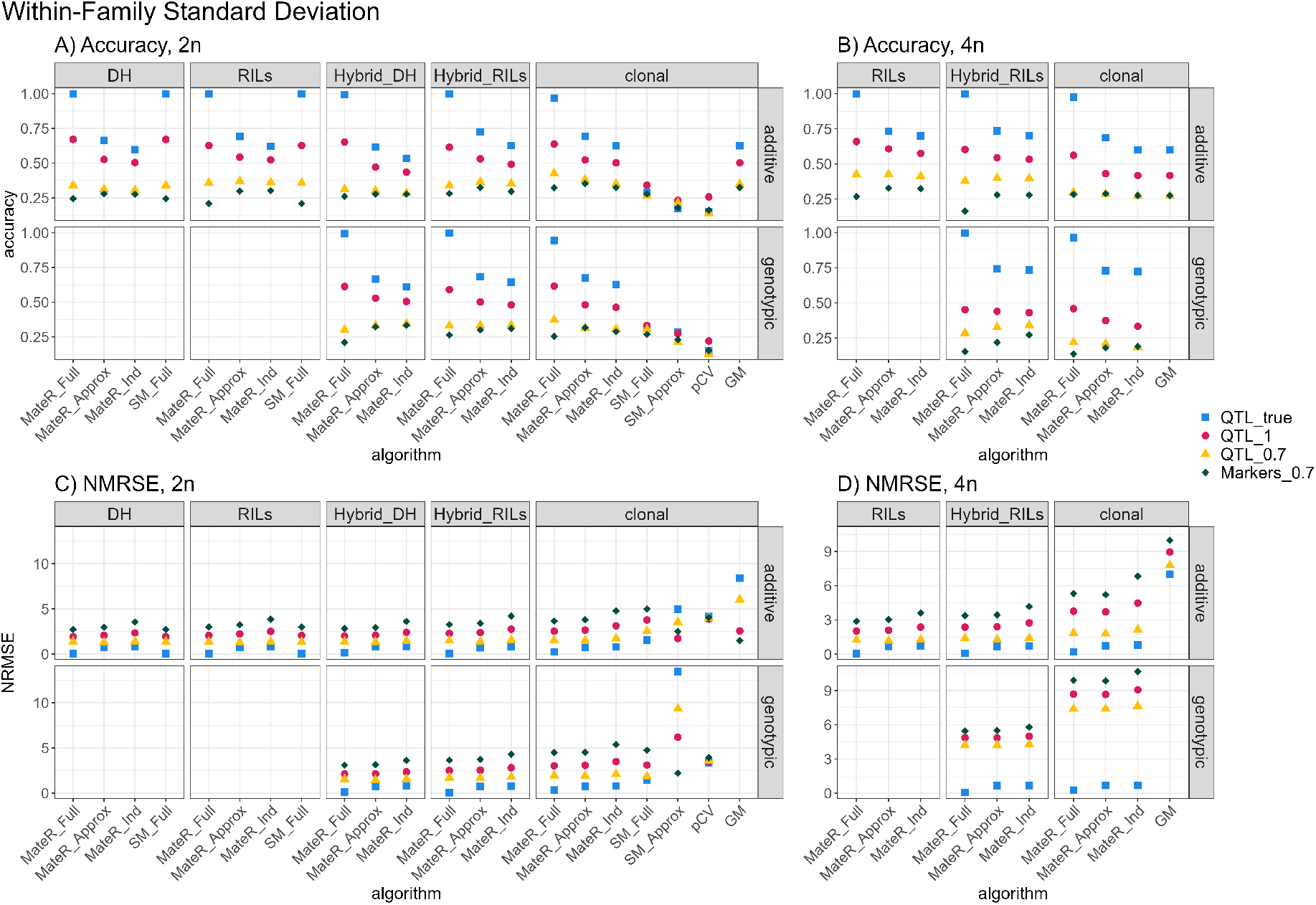
Within-family standard deviation. Agreement between predicted and true within-family SD across software packages. (A,B) Pearson correlation; (C,D) NRMSE. Panels show diploids (2n; A,C) and autotetraploids (4n; B,D). Within each panel, the top row uses additive values; the bottom row uses genotypic values (additive+dominance). Facet columns are population types (DH, RILs, Hybrid_DH, Hybrid_RILs, clonal). The *x*-axis lists the algorithm (MateR_Full, MateR_Approx, MateR_Ind, SM_Full, SM_Approx, pCV, GM). Colors/shapes encode how effects for prediction were obtained: 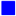 **QTL_true**; true QTL effects; 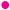 **QTL_1**; estimated QTL effects that exactly reproduce parental genome-wide genotypic values; 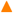 **QTL_0.7**; estimated QTL effects whose parental GEGVs correlate 0.7 with true genotypic values (TGVs); 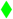 **Markers_0.7**; marker effects fitted on the 750 non-QTL loci giving parental GEGV–TGV correlation 0.7.

When phased genotypes were not available (`MateR_Approx`), accuracy dropped to around ~0.65–0.75 even with true QTL effects known (Figure 3A,B). Assuming marker independence (`MateR_Ind`) produced only a modest additional decrease and showed similar trends for NRMSE (Figure 3C,D). `GM` (Genomic mating) yielded identical accuracy to `MateR_Ind` but exhibited substantially higher NRMSE due to a coding error that doubles the predicted within-family standard deviation.

Degrading the quality of QTL or marker effects (`QTL_1`, `QTL_O.7`, `Markers_O.7`), reduced accuracy, increased NRMSE, and narrowed the advantage of `MateR_Full` over `MateR_Approx` and `MateR_Ind`. Under the most realistic setting, `Markers_O.7`, accuracy hovered around 0.25 and NRMSE frequently approached or exceeded 3, indicating systematic underestima-tion by more than three standard deviations. In this setting, `MateR_Approx` often showed slightly higher accuracy and comparable NRMSE to `MateR_Full`. `MateR_Ind` exhibited similar or marginally higher accuracy than `MateR_Approx` but with larger NRMSE Among non-MateR methods, `SM_Full` performed best and was comparable to `MateR_Full` in its three applicable scenarios (diploid *RILs, DH*, and *clonal*). In contrast, `SM_Approx` and `pCV`, usable only in the diploid *clonal* scenario, showed very low accuracy even with true QTL; `SM_Approx` also had an extremely large NRMSE, while `pCV` produced reasonable NRMSE but frequently predicted negative variances, which is problematic. Finally, `GM` applied only to additive values in clonal programs (both ploidies), matched `MateR_Ind` in accuracy, and consistently displayed much higher NRMSE.

### Proportion of standard deviation lost

Under random sampling, *PropSD* computed with Equation 15 was extremely accurate and presented an almost perfect correlation with the average true proportion of additive standard deviation lost, as shown in Figure 4. This was true for both diploid and autotetraploid crops.

**Figure 4.**
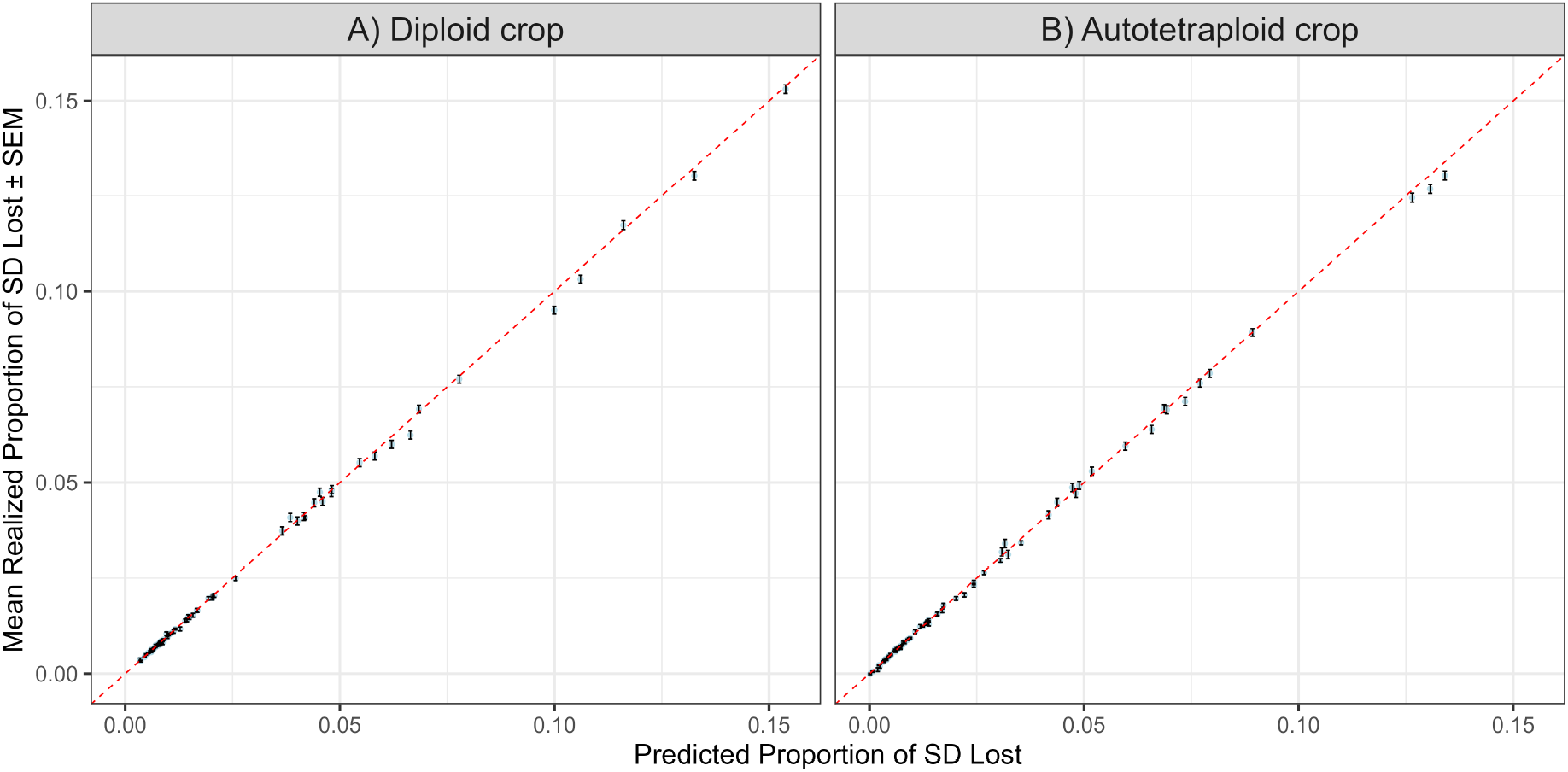
Correspondence between the *predicted* (horizontal axis) and the *true* (vertical axis) *PropSD* for diploid (A) and autotetraploid (B) datasets under **random sampling**. Each point belongs to a different random sample of selected parental lines and and shows the average across 10^5^ sets of random marker effects. Error bars indicate the standard error of the mean (SEM). The dashed red line marks perfect correspondence (identity).

Under selective pressure, the true proportion of genetic standard deviation lost was consistently and significantly larger than the predicted one (Figure 5A,D). By contrast, the predicted *PropSD* aligned much more closely with the realized loss of *genic* standard deviation (Figure 5B,D). In diploids, the realized genic loss tended to be lower than the predicted *PropSD*, whereas in autotetraploids the two were similar on average. Furthermore, genic standard deviation (Figure 5B,D) presented much lower dispersion than genetic standard deviation, as evidenced by the narrower boxplots (Figure 5B,D vs. A,C).

**Figure 5.**
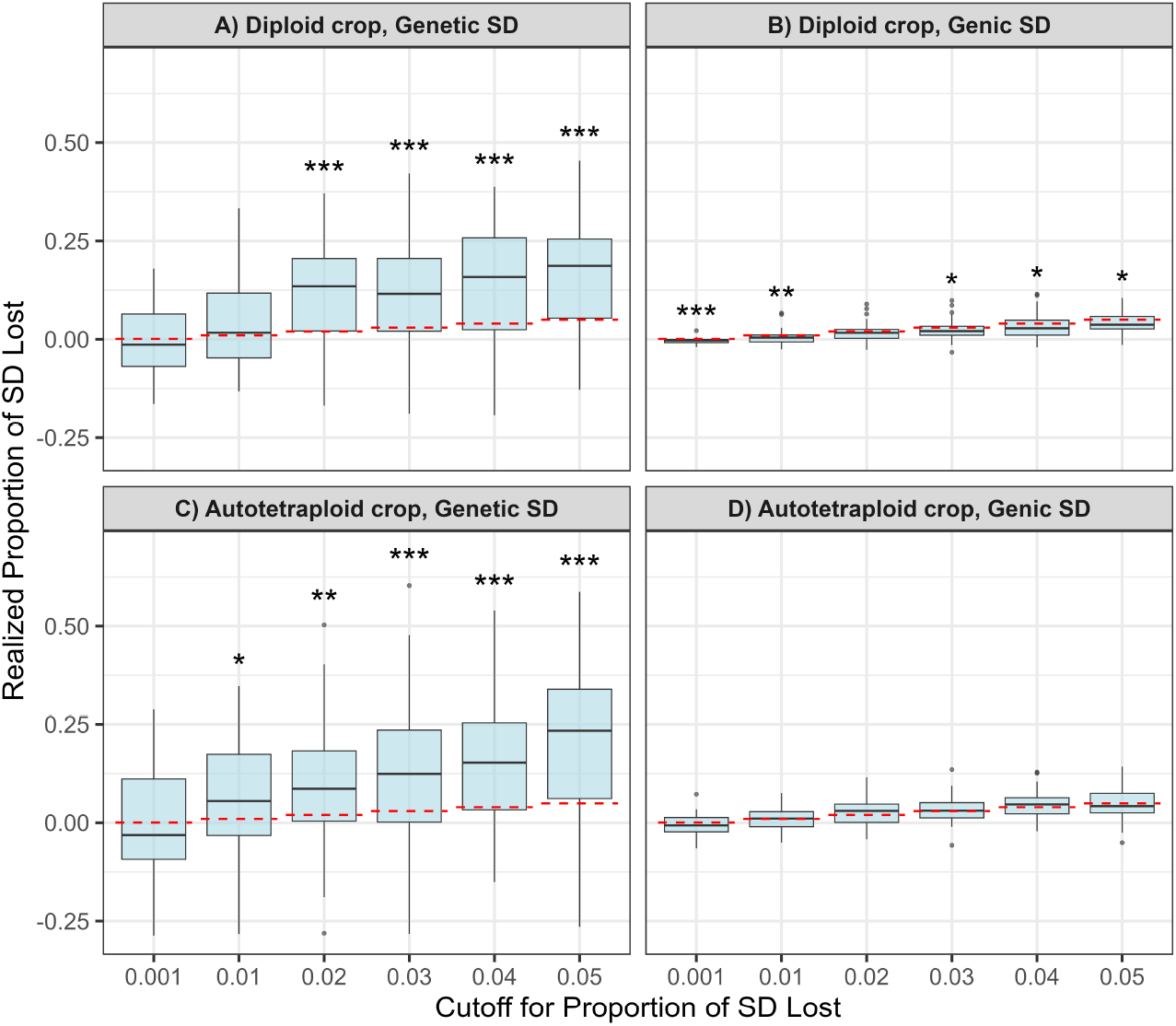
Correspondence between predicted and true *PropSD* under *selection* for diploid (A, B) and autotetraploid (C, D) datasets. Horizontal dashed red lines indicate the average *predicted PropSD* for the selected parents. Boxplots show the *realized* proportion of (A, C) *genetic* standard deviation lost (includes LD) and (B, D) *genic* standard deviation lost (LD-free). Asterisks denote the significance of paired Wilcoxon tests comparing predicted and realized values (0.01 < *p* < 0.05, 0.001 < *p* < 0.01, *p <* 0.001). Parametric tests were avoided because their assumptions were not met

## Discussion

In this section, we will consider the impact of the equations we have developed and other general considerations about the MateR package.

### Family mean

When only additive effects are considered, predicting family mean is trivial, as it corresponds to the mid-parental values. Kinghorn (2011); Akdemir and Sánchez (2016); Wolfe *et al*. (2021); Peixoto *et al*. (2025) use this approach and reach perfect accuracy when the true QTL effects are known, as seen in Figure 2. Our approach (Equation 4) is equivalent and obtains the same results for any set of QTL or marker effects.

When dominance effects are included, existing software relies on Equation 2 (Wolfe *et al*. 2021; Peixoto *et al*. 2025). This equation is limited to F1 generations and requires the genotypic parametrization, which limits its applicability and makes our approach superior. Furthermore, Figure 2 shows that it is not fully accurate even when the true QTL effects are known, which was unexpected. We further investigated this and found that Equation 2 outputs wrong results, as showcased in Supplementary File 1.

Finally, it is important to note that the accuracy of family mean prediction is very robust to QTL misidentification. In Figure 2, the QTL_true scenario was extremely similar to QTL_1, and QTL_0.7 was almost identical to Markers_0.7. This supports the notion that overall genome-wide prediction accuracy is the main factor influencing family mean accuracy, while knowing the distribution of the loci across the genome is secondary.

### Within-family standard deviation

Predicting family within-family standard deviation is harder than predicting family mean because correlations among loci contribute to the variance. Exact prediction therefore requires phased parental haplotypes to compute LD. In our benchmarks, `MateR_Full`, which uses phased haplotypes, was the only approach that consistently achieved near-perfect accuracy; `SM_Full` performed similarly only when dominance was absent and the parents were fully homozygous (Figure 3). `SM_Full` (Peixoto *et al*. 2025) is based on the equations of Lehermeier *et al*. (2017b), and was robust in the scenarios in which they are applicable. Our equations achieve comparable performance while also generalizing to dominance and autopolyploids.

Concerning the no-LD implementations, it is important to note that `GM` and `MateR_Ind` produced equivalent SD predictions except for a coding bug in `GM` that inflated SD by a factor of two. This did not change correlation-based accuracy but increased the normalized root mean square error (NRMSE).

Accuracy depended strongly on the fidelity of the effect map. Perfect accuracy was attainable only when the true QTL were known; it dropped markedly even in the *QTL_1* scenario, where causal effects were redistributed but genome-wide prediction accuracy remained ideal. This sensitivity follows directly from Eqs. 6 and 7: within-family variance aggregates cross-locus covariances, with weights proportional to locus effects, so the genomic placement of effects is decisive. In the most realistic setting (*Markers_0*.*7*), the accuracy for SD prediction was typically ≈0.25, which helps explain why the usefulness criterion can underperform mean-based selection in some studies (Wang *et al*. 2025). Nevertheless, modeling SD with accuracy ≈0.25 should be preferable to ignoring it, consistent with positive results reported elsewhere (Lehermeier *et al*. 2017b).

Overall, our results indicate that the current bottleneck for variance prediction is not the formulae but the availability of realistic marker-effect landscapes for quantitative traits (Rice *et al*. 2008; Korte and Farlow 2013; Ramstein *et al*. 2019; Clauw *et al*. 2024).

### Proportion of standard deviation lost

Balancing short-term gain with long-term diversity benefits from an interpretable metric. Because the additive standard deviation *σ*_*a*_ scales the expected response to selection, a given per-cycle value of PropSD implies (all else equal) that the next-cycle response satisfies *R*_*t*+1_ = (1 *−* PropSD)*R*_*t*_. This framing enables explicit, quantitative evaluation of long-term consequences for any chosen cutoff on PropSD. For such use to be reliable, predicted PropSD must track the realized loss of genetic variability. Under random sampling, the predicted PropSD matched the observed average across replicates (Figure 4), validating Eq. 14.

Under selection, however, the reduction in genetic SD exceeded predictions (Figure 5). This pattern is consistent with the Bulmer effect (Bulmer 1971): selection rapidly induces negative LD among QTL, depleting genetic variance beyond genic expectations. Our simulated base population lacked prior selection, so LD built up sharply in the first cycle. In contrast, long-running breeding programs are typically nearer Bulmer equilibrium, so deviations from Eq. 14 should be smaller or non-existent. Supporting this interpretation, the observed loss of *genic* SD (unaffected by LD) was similar to, or slightly lower than, the predicted PropSD. Multi-cycle evaluations will be useful to confirm the long-term correspondence between predicted PropSD and realized losses in additive SD.

### General considerations

A detailed comparison of existing genomic mating software with `MateR` is provided in Supplementary File 2. A key strength of `MateR` is its versatility. It accommodates diverse breeding schemes, including hybrid pipelines. In reciprocal recurrent selection, it identifies the optimal mating plan within a heterotic pool to maximize hybrid performance when progeny are testcrossed to the opposing pool, functionality that, to our knowledge, is unique.

`MateR` also optimizes cross replication (how many times a given pair of parents is crossed). Replicating a cross increases family size, thereby increasing within-family selection intensity and, potentially usefulness. Crucially, the benefit scales with within-family variance: if a family has (near) zero variance, additional sampling yields negligible gain. Hence, decisions about cross replication should be coupled to reliable variance prediction, as developed above.

## Conclusions

We develop a unified set of equations that exactly predict family mean and within-family variance when casual QTL effects are known across breeding schemes (including hybrid), with or without dominance, and for diploids, allopolyploids modeled as diploids, and autotetraploids. These results establish a coherent framework for expectation and dispersion that is parametrization consistent and able to consider selfing cycles and double haploids.

Empirically, predicting variance proved sensitive to the genomic placement of effects via LD, so accuracy declined under estimated marker effects; nevertheless, it rarely fell below *≈* 0.25 in realistic settings. `MateR_Approx` slightly outperformed `MateR_Full` under such conditions while being simpler, faster, and not requiring phased genotypes. We therefore recommend `MateR_Approx` as the default, reserving `MateR_Full` for cases with high-quality phased haplotypes and accurately estimated marker effects.

We also introduced *PropSD* as an interpretable metric for diversity management: because *σ*_*a*_ scales response to selection, a per-cycle loss *PropSD* implies *R*_*t*+1_ = (1 *− PropSD*)*R*_*t*_ (all else equal). This metric makes explicit the trade-off between short-term gain and long-term potential, enabling principled tuning of mating plans. Together, these advances render genomic mating operational at scale, while clarifying the main bottleneck, accurate, LD-aware effect estimation, and the levers available to optimize long-term genetic progress.

## Supporting information

Supplementary Files 1 and 2. Description available in manuscript

## Data availability

The code used to generate results and minimal input data to reproduce the figures will be made publicly available upon publication at GITHUB. Any additional resources are provided in Files S1–S2.

## Supplementary Materials

- Supplementary File 1: R script providing reproducible counterexamples showing that Eq. 2 does not hold across the tested scenarios (several F1 crosses under explicit dominance), with detailed comments and expected outputs.
- Supplementary File 2: Excel workbook offering a feature-by-feature comparison of genomic mating software, including objective functions, parametrizations (additive/dominance), generation scope, polyploid support, input requirements (markers vs. phased haplotypes), and optimization capabilities.

## Author contribution statement

JFG conducted the mathematical development, developed `MateR` package, and performed the statistical analyses. SMM developed the transition-matrix framework for predicting genotypic frequencies after any number of selfing cycles and tested MateR under many scenarios. JIyS conceived the study and played a role in study conceptualization, coordination, manuscript drafting, and securing funding. JFG and SMM played a crucial role in software/equation testing and iterative improvements. All authors jointly interpreted the results and co-wrote the manuscript.

## Appendices Appendix 1: Genotypic frequencies and extension to polyploids

### Diploid crop, prediction of genotypic frequencies

In a diploid crop, for a single gene with two alleles, *A* and *a*, there are three possible genotypes: *AA, Aa*, and *aa*. When we cross two parents, we want to predict the frequencies of each genotype (*P*_*AA*_, *P*_*Aa*_, *P*_*aa*_) in their offspring family. These frequencies can be obtained from basic Mendelian genetics. We also need to extend them for self-pollination (selfing) over multiple generations. Selfing changes the genotypic frequencies in a predictable way:

- The heterozygous genotype *Aa* becomes less frequent by half each generation.
- Each homozygous genotype (*AA* and *aa*) increases by one quarter of the *Aa* frequency from the previous generation.

Mathematically:

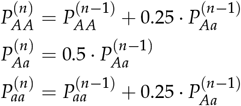

After rearranging and simplifying, we can generalize the genotype frequencies after *n* generations of selfing for different parental combinations, as shown in Table 2.

**Table 2.** Genotypic frequencies in a diploid locus with alleles *A* and *a*, for different parental combinations after *n* cycles of selfing. The first two columns contain the genotypes of the two parents. The other columns contain the frequencies of each possible genotype.

| P1 | P2 | AA (n=0) | Aa (n=0) | aa (n=0) | AA (any $n$) | Aa (any $n$) | aa (any $n$) |
| --- | --- | --- | --- | --- | --- | --- | --- |
| AA | AA | 1 | 0 | 0 | 1 | 0 | 0 |
| aa | aa | 0 | 0 | 1 | 0 | 0 | 1 |
| AA | aa | 0 | 1 | 0 | $\frac{1}{2} - \left(\frac{1}{2}\right)^{n+1}$ | $\left(\frac{1}{2}\right)^n$ | $\frac{1}{2} - \left(\frac{1}{2}\right)^{n+1}$ |
| Aa | Aa | $\frac{1}{4}$ | $\frac{1}{2}$ | $\frac{1}{4}$ | $\frac{1}{2} - \left(\frac{1}{2}\right)^{n+2}$ | $\left(\frac{1}{2}\right)^{n+1}$ | $\frac{1}{2} - \left(\frac{1}{2}\right)^{n+2}$ |
| AA | Aa | $\frac{1}{2}$ | $\frac{1}{2}$ | 0 | $\frac{3}{4} - \left(\frac{1}{2}\right)^{n+2}$ | $\left(\frac{1}{2}\right)^{n+1}$ | $\frac{1}{4} - \left(\frac{1}{2}\right)^{n+2}$ |
| Aa | aa | 0 | $\frac{1}{2}$ | $\frac{1}{2}$ | $\frac{1}{4} - \left(\frac{1}{2}\right)^{n+2}$ | $\left(\frac{1}{2}\right)^{n+1}$ | $\frac{3}{4} - \left(\frac{1}{2}\right)^{n+2}$ |

It is important to note that allopolyploids are normally modeled as diploids. Therefore, Table 2 can also be applied to them.

### Extension to autotetraploids

The equations used for diploid crops can be extended to autopolyploid species, such as autotetraploids, by adapting the genotype states and their expected values. In the case of autotetraploids, we assume **bivalent and non-preferential pairing** during meiosis.

At a biallelic locus *j* in an autotetraploid family *F*, there are five possible genotypes: *AAAA, AAAa, AAaa, Aaaa*, and *aaaa*. The corresponding genotypic frequencies are denoted as:

- *PAAAA*_*F,j*_ for genotype *AAAA*
- *PAAAa*_*F,j*_ for genotype *AAAa*
- *PAAaa*_*F,j*_ for genotype *AAaa*
- *PAaaa*_*F,j*_ for genotype *Aaaa*
- *Paaaa*_*F,j*_ for genotype *aaaa*

To translate these genotypes into numeric form, we use two types of incidence matrices:

- **Additive incidence matrix:** represents the number of copies of the *a* allele. *AAAA* = 0, *AAAa* = 1, *AAaa* = 2, *Aaaa* = 3, and *aaaa* = 4
- **Dominance incidence matrix:** represents the presence of heterozygosity (non-homozygosity). *AAAa* = 3, *AAaa* = 4, *Aaaa* = 3, while *AAAA* = 0 and *aaaa* = 0. This can be generalized to any ploidy as *x*(*ϕ− x*), where *ϕ* is the degree of ploidy and *x* refers to the additive score, i.e., the number of times the *A* allele is present Labroo *et al*. (2023). This equation only captures the digenic dominance component, and discards trigenic or higher-order interactions, which is a common simplification Endelman *et al*. (2018).

These incidence scores belong to the genotypic parametrization. In the breeding parametrization, for a minor allele frequency in the reference population *p*, we would need to subtract *ϕp* from the additive scores and use 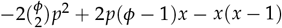 for the dominance scores (Endelman 2023).

Continuing with the genotypic parametrization for the sake of simplicity, and using the same nomenclature as in Equations 3 and 5, we define the expected genotypic value (*µ*_*F,j*_) and variance 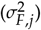 at locus *j* for family *F* as follows:

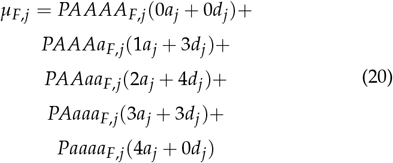

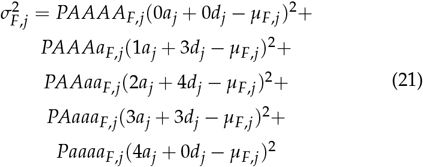

Equations 4, 7, and 13 remain valid for autotetraploids. The only requirement is to compute the locus-specific means and variances using Equations 20 and 21, which account for the additional genotypic classes and dosage effects in tetraploid contexts. These per-locus values are then summed or combined across the genome as in the diploid case.

### Autotetraploid Crop, Prediction of Genotypic Frequencies

The genotypic frequencies of the F1 generation (i.e., with no cycles of selfing, *n* = 0) in autotetraploid crops are summarized in Table 3. These initial frequencies form the basis for computing the genotypic frequencies in subsequent generations after *n* cycles of selfing. The frequencies from Table 3 are derived from the convolution of two hypergeometric distributions, which represent the probabilities of sampling alleles from the gametes of each parent under random bivalent pairing Serang *et al*. (2012); Gerard *et al*. (2018). While autotetraploids are the only autopolyploids currently supported by MateR, this methodology could be used to extend it to any degree of ploidy.

**Table 3.** Genotypic frequencies in an autotetraploid locus with two alleles (*A* and *a*), obtained when crossing two parents (P1 and P2). The first two columns show the parental genotypes. The next five columns display the frequencies of genotypes in the F1 generation, i.e., without any selfing.

| P1 | P2 | AAAA | AAAa | AAaa | Aaaa | aaaa |
| --- | --- | --- | --- | --- | --- | --- |
| AAAA | AAAA | 1 | 0 | 0 | 0 | 0 |
| aaaa | aaaa | 0 | 0 | 0 | 0 | 1 |
| AAAA | AAAa | $\frac{1}{2}$ | $\frac{1}{2}$ | 0 | 0 | 0 |
| Aaaa | aaaa | 0 | 0 | 0 | $\frac{1}{2}$ | $\frac{1}{2}$ |
| AAAA | AAaa | $\frac{6}{36}$ | $\frac{24}{36}$ | $\frac{6}{36}$ | 0 | 0 |
| AAaa | aaaa | 0 | 0 | $\frac{6}{36}$ | $\frac{24}{36}$ | $\frac{6}{36}$ |
| AAAA | Aaaa | 0 | $\frac{1}{2}$ | $\frac{1}{2}$ | 0 | 0 |
| AAAa | aaaa | 0 | 0 | $\frac{1}{2}$ | $\frac{1}{2}$ | 0 |
| AAAa | AAAa | $\frac{1}{4}$ | $\frac{1}{2}$ | $\frac{1}{4}$ | 0 | 0 |
| Aaaa | Aaaa | 0 | 0 | $\frac{1}{4}$ | $\frac{1}{2}$ | $\frac{1}{4}$ |
| AAAa | AAaa | $\frac{3}{36}$ | $\frac{15}{36}$ | $\frac{15}{36}$ | $\frac{3}{36}$ | 0 |
| AAaa | Aaaa | 0 | $\frac{3}{36}$ | $\frac{15}{36}$ | $\frac{15}{36}$ | $\frac{3}{36}$ |
| AAAa | Aaaa | 0 | $\frac{1}{4}$ | $\frac{1}{2}$ | $\frac{1}{4}$ | 0 |
| AAaa | AAaa | $\frac{1}{36}$ | $\frac{8}{36}$ | $\frac{18}{36}$ | $\frac{8}{36}$ | $\frac{1}{36}$ |
| AAAA | aaaa | 0 | 0 | 1 | 0 | 0 |

To compute genotypic frequencies after selfing, we use a selfing transition matrix (*X*), which has the values from Table 4 in autotetraploids. The frequencies in *X* are particular cases of Table 3. *X* is a column-stochastic square matrix where each parental genotype class is acting like a “state” in a Markov-chain. Its columns contain the conditional probabilities of transitioning from parental to progeny states, with “time” being the number of selfing cycles. For any parental genotype *j* and any offpsring genotype *i*:

**Table 4.** *X* matrix in table format. It contains the parental genotypes in the columns, and each row contains the probability of an offspring genotype after selfing the parent.

|  | AAAA | AAAa | AAaa | Aaaa | aaaa |
| --- | --- | --- | --- | --- | --- |
| AAAA | 1 | $\frac{1}{4}$ | $\frac{1}{36}$ | 0 | 0 |
| AAAa | 0 | $\frac{1}{2}$ | $\frac{8}{36}$ | 0 | 0 |
| AAaa | 0 | $\frac{1}{4}$ | $\frac{18}{36}$ | $\frac{1}{4}$ | 0 |
| Aaaa | 0 | 0 | $\frac{8}{36}$ | $\frac{1}{2}$ | 0 |
| aaaa | 0 | 0 | $\frac{1}{36}$ | $\frac{1}{4}$ | 1 |

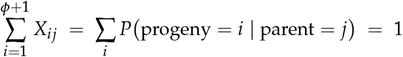

Where *ϕ* is the degree of ploidy and the number of possible genotypic classes is therefore *ϕ* + 1. Based on the law of total probability, the overall probability that an offspring ends up being of genotype *i* has to be:

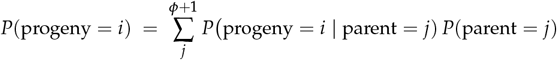

For any family *F*, this can be generalized in matrix notation as follows:

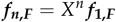

where *f*_1,*F*_ is a vector of F1 progeny genotypic frequencies and *f*_*n,F*_ is a vector of progeny genotypic frequencies after *n* selfing cycles.

## Appendix 2: Extension of family mean and variances to multi-trait

The equations described before for computing genotypic means and variances at the single-locus level can be readily extended to a multi-trait context. In multi-trait selection, the goal is often to combine information from multiple traits into a single value that reflects overall genotypic value or fitness. This is commonly achieved using a **selection index**.

Let:

- *M* be the *n* × *p* matrix of additive marker scores, where *n* is the number of individuals and *p* is the number of markers.
- *W* be the corresponding *n* × *p* matrix of dominance scores, using the same codification as previously described.
- ***a***_***t***_ and ***d***_***t***_ be *p ×* 1 vectors of additive and dominance effects, for trait *t*.
- *c*_*t*_ be a scalar weight (coefficient derived from economic value or importance) assigned to trait *t*, used to combine traits into a single selection index.
- *n*_*t*_ be the number of traits included in the index.

The genotypic values for individuals with respect to trait *t* are given by:

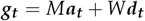

The multi-trait selection index scores (***IS***) for all individuals is computed as a weighted linear combination of their genotypic values across traits:

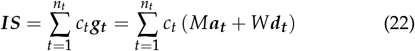

Using the distributive property of matrix multiplication, this expression simplifies to:

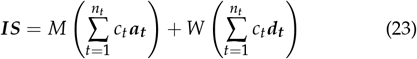

We can then define the **multi-trait additive and dominance marker effects** as:

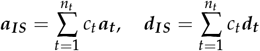

Thus, the multi-trait selection index is:

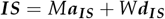

This representation reduces the multi-trait selection problem to a **single-trait framework**, where the trait is the index itself. Consequently, we can use the previously defined equations for genotypic mean (Equation 3) and variance (Equation 5) by replacing *a*_*j*_ and *d*_*j*_ with 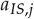 and 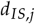.

## Appendix 3: Mathematical derivation for the proportion of additive standard deviation lost

Let 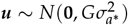 be the additive values for the entire parental pool and 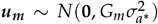 be the additive values for the selected parents. It is important to note that 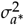 is a scaling factor (often estimated with REML) but it has no biological meaning due to the improper scaling of the *G* matrix Fernández-González and Isidro y Sánchez (2025). We will use an asterisk to identify the non-interpretable scaling factors, while variance components with actual biological meaning will be written without the asterisk.

Due to selection, the genetic standard deviation in *a*_*m*_ is often lower than that of *a*. The proportion of standard deviation retained after selection is 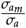. Thus, the proportion of standard deviation lost (*PropSD*) is 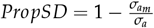. The actual value of the standard deviations in the parental pool (*σ*_*a*_) and in the selected parents 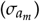 are unknown, but it is possible to estimate them as the expected standard deviation of an infinite amount of samples from the corresponding multivariate distributions: 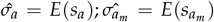. To obtain these estimators, we will first obtain the expected sample variance:

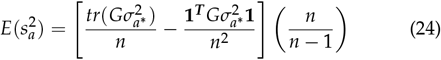

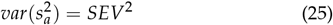

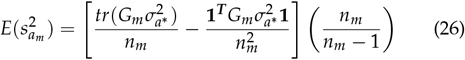

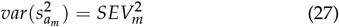

Where *n* is the total number of parents, *n*_*m*_ is the number of parents selected for the mating plan, *SEV* is the standard error of the variance computed as in Fernández-González and Isidro y Sánchez (2025) from the distribution 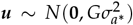 and *SEV*_*m*_ is the standard error of the variance corresponding to 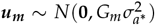.It is important to note that equations 24 and 26 are computed following Searle *et al*. (1992); Endelman (2023) with the addition of the correction factor 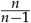 to account for the fact that we are computing the expectation of sample variance, not population variance. Thus, a degree of freedom has to be expended in the computation of the sample mean before calculating the variance.

Additionally, while it can be tempting to estimate 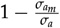 as 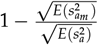, this is not possible due to the fact that 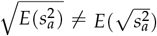. Thus, we need to find a way for computing 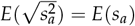. Thankfully, it is known that sample variance follows a gamma distribution. Therefore:

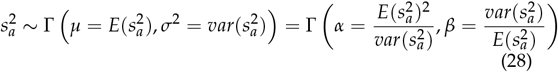

Where *α* and *β* are the shape and scale parameters of a gamma distribution. To compute *E*(*s*_*a*_) we can use the fact that the square root of a gamma distributed variable is known to follow a Nakagami distribution. Similarly, we can use the probability density function of the gamma distribution and compute *E*(*s*_*a*_) as the moment of order 1/2. The probability density function of a gamma distribution is:

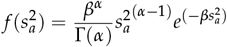

Where Γ(*·*) refers to the gamma function. 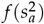 is defined for all 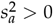. Gamma distributions have the property that Γ(*α, β*) = *β ·* Γ(*α*, 1). Thus, if *β* is taken out of the gamma distribution, we can simplify the probability density function to:

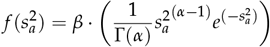

Therefore, we can define 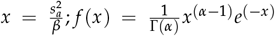.

Next, applying the equation for the moment of order 1/2 of a continuous distribution:

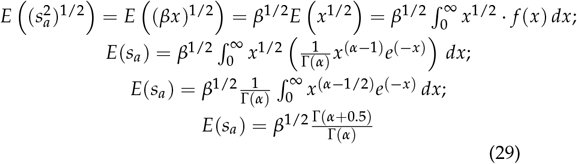

Where *α* and *β* are computed as described in equations 24, 25 and 28. The same equation can be applied to the computation of 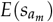. Therefore, we can calculate the proportion of standard deviation lost as:

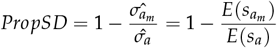

It is important to note that both *E*(*s*_*a*_) and 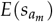 depend solely on 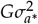 and 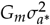. If the *G* matrix is multiplied by a scalar, it holds that *E*(*s*_*a*_) gets multiplied by the square root of the same scalar. Therefore:

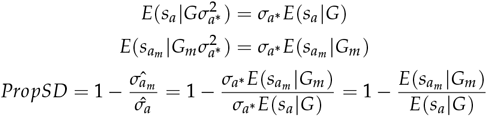

As a result, the REML estimation of 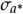 is not required for the computation of *PropSD*, only *G* and *G*_*m*_ are needed.

## Appendix 4. Inbreeding rate

Genetic diversity is often conceptualized in terms of inbreeding (*F*) for genomic mating Kinghorn (2011); Akdemir and Sánchez (2016); Gorjanc and Hickey (2018); Endelman (2025). However, while inbreeding can be computed accurately from a numerator relationship matrix (*A*) if a good pedigree is available (which is rare), it presents some error when computed from genomic relationships Wang (2014); Endelman (2024). The likely reasons behind this are that: i) the *G* matrix captures identity by state, not only identity by descent and ii) the scaling of the *G* matrix is often wrong, as the VanRaden method relies on very simplistic and unrealistic assumptions Fernández-González and Isidro y Sánchez (2025). Thus, we recommend using *PropSD* instead. However, if desired, MateR also supports controlling diversity with inbreeding rate (Δ*F*). Δ*F* is computed as Woolliams *et al*. (2015); Endelman (2025):

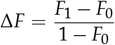

Where *F*_0_ is the inbreeding value in the parental pool and *F*_1_ is the inbreeding value in the selected crosses. We can compute *F*_0_ and *F*_1_ as the average inbreeding of all genotypes in the corresponding generation. Inbreeding is often computed as the diagonal of the relationship matrix minus one, but this can be very problematic in some scenarios. For instance, for dobule haploids, all diagonal elements in *A* are always 2. Therefore, the average inbreeding for any population would be 1 regardless of how related the genotypes are to each other, i.e., we are not capturing group coancestry. Therefore, we need an alternative. According to Xu (2022), the inbreeding of an individual is equal to the average coancestry of its two parents, which itself is half their relationship in *A*. Therefore, *F*_1_ can be computed as the mean of the off-diagonal elements of the *A* matrix for their parents divided by two:

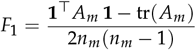

Where *A*_*m*_ is the numerator relationship matrix for the parents of the mating plan, with each parent replicated as many times as the number of crosses in which it is involved. *n*_*m*_ is the number of rows or columns in *A*_*m*_.

For *F*_0_ it seems that we would need the *A* matrix for the previous generation, which is often unavailable. However, we know that, for four parents *a, b, c, d* and two offspring *ab* and *cd* obtained by crossing the respective parents, the relationship between them is Akdemir and Sánchez (2016):

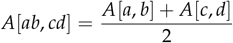

Thus, *F*_0_ can be computed as:

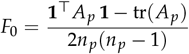

Where *A*_*p*_ is the relationship matrix for the parental pool and *n*_*p*_ is the number of parents.

If the *G* matrix is used instead, we can simply compute inbreeding as:

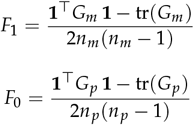

It is very important to use the VanRaden VanRaden (2008) relationship matrix in this context, as it provides a scaling of *G* that makes it analogous to *A*. Furthermore, this method of computing inbreeding is the same as the third method in Morales-González *et al*. (2020). However, as explained earlier, using the *G* matrix will likely incur some errors due to improper scaling and contamination from identity by state.

## Appendix 5. Standard error of the mean and selection intensity

To assess the diversity of a mating plan, we treat *P*_*m*_ as a sample from the entire parental population, which is assumed to have mean-centered additive values (i.e., mean zero). Using 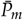 to denote the average additive value of the selected parents, we estimate the standard error of the mean (SEM) using the following expression, which accounts for correlations among samples Fernández-González and Isidro y Sánchez (2025):

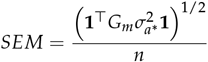

where *G*_*m*_ is the genomic relationship matrix of the parents selected for the mating plan, **1** is a vector of ones, and *n* is the number of rows (or columns) in *G*_*m*_. SEM can be related to selection intensity so as to make it more interpretable. 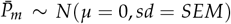 has zero mean because sample mean is an unbiased estimator. We scale 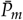 to standard deviation units by:

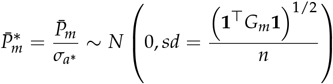

We assume that the overall parental average (*µ*) is zero due to mean-centered BLUPs. Therefore, it holds that 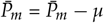, which can be interpreted as a selection differential. Similarly, 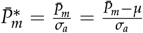can be viewed as a selection intensity. There-fore, we can directly compare 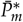 with the actual selection intensity of the selected parents (*i*_*realized*_), which can be calculated using the genomic estimated additive values of the parental genotypes. We are interested in the probability that the scaled average of the selected parents exceeds *i*_*realized*_:

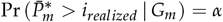

This can be rearranged as:

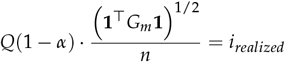

where *Q*(*x*) is the quantile function of the standard normal distribution. This allows to compute *α*, which is the likelihood of a solution with higher gain than the current mating plan with similar diversity.

MateR allows to control diversity loss using SEM as a metric directly. It also always computes *α* internally for any optimal mating plan, which allows to evaluate the goodness of the optimization. The lower the value of *α*, the more unlikely it is that a solution better than the current one exists. Thus, low *α* values indicate a good optimization quality.

It is important to note that *α* values are calculated based only on the additive genomic relationships. Thus, they are unable to capture dominance and they become meaningless in the presence of heterosis.

## Literature cited

Akdemir D, Rio S, Isidro y Sánchez J. 2021a. Trainsel: an r package for selection of training populations. Frontiers in genetics. 12:655287.

Akdemir D, Rio S, Sánchez IY et al. 2021b. TrainSel: an R package for selection of training populations. Frontiers in Genetics. 12:607.

Akdemir D, Sánchez JI. 2016. Efficient breeding by genomic mating. Frontiers in Genetics. 7:1–12.

Bulmer M. 1971. The effect of selection on genetic variability. The American Naturalist. 105:201–211.

Clauw P, Ellis TJ, Liu HJ, Sasaki E. 2024. Beyond the standard gwas—a guide for plant biologists. Plant And Cell Physiology. p. pcae079.

Endelman JB. 2023. Fully efficient, two-stage analysis of multienvironment trials with directional dominance and multi-trait genomic selection. Theoretical and Applied Genetics. 136:65.

Endelman JB. 2024. Directional dominance in polyploids: Trait analysis and mate selection. In:. Vienna, Austria. Presented at a conference.

Endelman JB. 2025. Genomic prediction of heterosis, inbreeding control, and mate allocation in outbred diploid and tetraploid populations. Genetics. 229:iyae193.

Endelman JB, Carley CAS, Bethke PC, Coombs JJ, Clough ME, da Silva WL, De Jong WS, Douches DS, Frederick CM, Haynes KG et al. 2018. Genetic variance partitioning and genome-wide prediction with allele dosage information in autotetraploid potato. Genetics. 209:77–87.

Falconer D. 1989. Introduction to quantitative genetics. Longman Scientific & Technical. New York.

Fernández-González J, Isidro y Sánchez J. 2025. Maximizing the accuracy of genetic variance estimation and using a novel generalized effective sample size to improve simulations. Theoretical and Applied Genetics. 138:1–21.

Gerard D, Ferrão LFV, Garcia AAF, Stephens M. 2018. Genotyping polyploids from messy sequencing data. Genetics. 210:789–807.

Gorjanc G, Hickey JM. 2018. Alphamate: a program for optimizing selection, maintenance of diversity and mate allocation in breeding programs. Bioinformatics. 34:3408–3411.

Grundy B, Villanueva B, Woolliams J. 1998. Dynamic selection procedures for constrained inbreeding and their consequences for pedigree development. Genetics Research. 72:159–168.

Kinghorn BP. 2011. An algorithm for efficient constrained mate selection. Genetics Selection Evolution. 43:1–9.

Korte A, Farlow A. 2013. The advantages and limitations of trait analysis with gwas: a review. Plant methods. 9:1–9.

Labroo MR, Endelman JB, Gemenet DC, Werner CR, Gaynor RC, Covarrubias-Pazaran GE. 2023. Clonal diploid and autopolyploid breeding strategies to harness heterosis: insights from stochastic simulation. Theoretical and Applied Genetics. 136:147.

Lehermeier C, de Los Campos G, Wimmer V, Schön CC. 2017a. Genomic variance estimates: With or without disequilibrium covariances? Journal of Animal Breeding and Genetics. 134:232–241.

Lehermeier C, Teyssèdre S, Schön CC. 2017b. Genetic gain increases by applying the usefulness criterion with improved variance prediction in selection of crosses. Genetics. 207:1651–1661.

Lynch M, Walsh B et al. 1998. Genetics and analysis of quantitative traits. volume 1. Sinauer Sunderland, MA.

Meuwissen T. 1997. Maximizing the response of selection with a predefined rate of inbreeding. Journal of animal science. 75:934–940.

Meuwissen THE, Hayes BJ, Goddard ME. 2001. Prediction of total genetic value using genome-wide dense marker maps. Genetics. 157:1819–1829.

Morales-González E, Saura M, Fernández A, Fernández J, Pong-Wong R, Cabaleiro S, Martínez P, Martín-García A, Villanueva B. 2020. Evaluating different genomic coancestry matrices for managing genetic variability in turbot. Aquaculture. 520:734985.

Peixoto MA, Amadeu RR, Bhering LL, Ferrão LFV, Munoz PR, Resende Jr MF. 2025. Simplemating: R-package for prediction and optimization of breeding crosses using genomic selection. The Plant Genome. 18:e20533.

Ramstein GP, Jensen SE, Buckler ES. 2019. Breaking the curse of dimensionality to identify causal variants in Breeding 4. Theoretical and Applied Genetics. 132:559–567.

Rice TK, Schork NJ, Rao D. 2008. Methods for handling multiple testing. Advances in genetics. 60:293–308.

Schnell F, Utz H. 1975. Bericht über die arbeitstagung der vereinigung österreichischer pflanzenzüchter. Gumpenstein: BAL Gumpenstein. pp. 243–8.

Searle SR, Casella G, McCulloch CE. 1992. Variance components. John Wiley & Sons.

Serang O, Mollinari M, Garcia AAF. 2012. Efficient exact maximum a posteriori computation for bayesian snp genotyping in polyploids. PLoS ONE. 7:e30906.

Toro M, Perez-Enciso M. 1990. Optimization of selection response under restricted inbreeding. Genetics Selection Evolution. 22:93–107.

VanRaden P. 2008. Efficient methods to compute genomic predictions. Journal of Dairy Science. 91:4414–4423.

Vitezica ZG, Varona L, Legarra A. 2013. On the additive and dominant variance and covariance of individuals within the genomic selection scope. Genetics. 195:1223–1230.

Wang F, Feldmann MJ, Runcie DE. 2025. Why usefulness is rarely useful. G3: Genes, Genomes, Genetics. 15:jkae296.

Wang J. 2014. Marker-based estimates of relatedness and inbreeding coefficients: an assessment of current methods. Journal of Evolutionary Biology. 27:518–530.

Wolfe MD, Chan AW, Kulakow P, Rabbi I, Jannink JL. 2021. Genomic mating in outbred species: predicting cross usefulness with additive and total genetic covariance matrices. Genetics. 219:iyab122.

Woolliams JA, Berg P, Dagnachew BS, Meuwissen T. 2015. Genetic contributions and their optimization. Journal of Animal Breeding and Genetics. 132:89–99.

Wray N, Goddard M. 1994. Increasing long-term response to selection. Genetics Selection Evolution. 26:431–451.

Wray NR, Thompson R. 1990. Prediction of rates of inbreeding in selected populations. Genetics Research. 55:41–54.

Wright S. 1934. The method of path coefficients. The annals of mathematical statistics. 5:161–215.

Xu S. 2022. Quantitative genetics. Springer.

Zhong S, Jannink JL. 2007. Using quantitative trait loci results to discriminate among crosses on the basis of their progeny mean and variance. Genetics. 177:567–576.

