## Supplementary Files 1 and 2. Description available in manuscript for "MateR: a novel framework for computing the usefulness criterion and applying genomic mating": Supplementary_File_1.R

```
a = 2 #additive marker effect (substitution effect)
d = 1 #dominance effect

################################################################################
#Fully homozygous parents for different alleles
################################################################################

genoP1 = 0
genoP2 = 2
p1 = genoP1/2
q1 = 1-p1
y = (genoP1-genoP2)/2


#This line of code only works for fully homozygous parents that generate a uniform F1!
genoF1 = (genoP1+genoP2)/2

#True value
(MeanF1 = genoF1*a + (genoF1==1)*d)

#Falconer formula (Equation 14.6 from the 1996 edition)
(FalconerMeanF1 = a*(p1-q1-y)+d*(2*p1*q1 + y*(p1-q1)))

#The true value and the predicted one do not match


################################################################################
#Fully homozygous parents for same allele
################################################################################

genoP1 = 0
genoP2 = 0
p1 = genoP1/2
q1 = 1-p1
y = (genoP1-genoP2)/2


#This only works for fully homozygous parents that generate a uniform F1!
genoF1 = (genoP1+genoP2)/2

#True value
(MeanF1 = genoF1*a + (genoF1==1)*d)

#Falconer formula (Equation 14.6 from the 1996 edition)
(FalconerMeanF1 = a*(p1-q1-y)+d*(2*p1*q1 + y*(p1-q1)))

#The true value and the predicted one do not match


################################################################################
#Fully homozygous parents for same allele
################################################################################

genoP1 = 2
genoP2 = 2
p1 = genoP1/2
q1 = 1-p1
y = (genoP1-genoP2)/2


#This only works for fully homozygous parents that generate a uniform F1!
genoF1 = (genoP1+genoP2)/2

#True value
(MeanF1 = genoF1*a + (genoF1==1)*d)

#Falconer formula (Equation 14.6 from the 1996 edition)
(FalconerMeanF1 = a*(p1-q1-y)+d*(2*p1*q1 + y*(p1-q1)))

#The true value and the predicted one do not match
```
